# A neural circuit for rapid control of hunger by arousal

**DOI:** 10.1101/2025.03.07.642010

**Authors:** Seher Kosar, Eva Bracey, Yihui Du, Daria Peleg-Raibstein, Denis Burdakov

**Affiliations:** ETH Zürich: Department of Health Sciences and Technology (D-HEST); Neuroscience Center Zürich (ZNZ); Institute of Food, Nutrition and Health; Institute for Neuroscience. Schwerzenbach, Switzerland

## Abstract

The brain must constantly integrate internal needs with external cues, requiring it to evaluate potential rewards and prioritize urgent environmental demands. How arousal systems control hunger to achieve this is a fundamental question. We targeted the functional connection from arousal hypocretin/orexin neurons (HONs) to hunger (AGRP) neurons and discovered that it operates through a local GABAergic relay, revealing a circuit for rapid, arousal-driven hunger suppression. We found this suppression deployed in several critical functions. In appetitive evaluation, it controls food value perception by suppressing AGRP based on a food’s predicted caloric worth. In adaptive vigilance, it creates an opportunity to prioritize other needs, such as responding to novel, arousing stimuli. During spontaneous pupil dilations, it creates ‘hunger-free’ windows for optimal assessment of the environment. These findings reveal a fundamental inhibitory circuit that allows pupil-linked arousal to dynamically gate hunger on a moment-to-moment basis, providing a new framework for how the brain balances competing internal and external demands.

## Introduction

The brain must constantly integrate internal needs with external cues, a process critical for survival. For a foraging animal, this presents a fundamental challenge: the potent drive to eat, governed by primary hunger circuits[1–3], must be dynamically modulated to evaluate both opportunities and dangers[4]. This modulation serves two distinct and critical purposes. First, it must evaluate appetitive cues of a potential meal, a computation necessary to assess the predicted value of a food source[1, 5–8]. Second, more urgently, it must instantly suppress aversive hunger[7], allowing attention to be reallocated to an unexpected external stimulus, such as a sudden rustle in the undergrowth. This rapid de-prioritization is part of what Pavlov called the ‘What is that?’ response[9], an ancient arousal adaptation that enables an organism to halt ongoing behaviors to assess and respond to potentially important events [10–12]. How the brain’s arousal systems exert such flexible, context-dependent control over a core homeostatic drive has been a major unresolved question [1].

Two evolutionarily-conserved neural populations may be involved in this interplay between arousal and hunger. One is the hypocretin/orexin neurons (HONs) in the lateral hypothalamus, which are rapidly engaged by external stimuli, both food cues and unexpected environmental cues [13–20]. HON activity broadcasts arousal information to multiple brain regions, and promotes arousal [21–27]. The other is agouti-related peptide/neuropeptide Y co-expressing neurons (AGRP neurons) in the hypothalamic arcuate nucleus, which are thought to be a key cause of hunger, and are activated by energy depletion [1, 28–32]. AGRP neurons are inhibited by multiple aspects of feeding: rapid feedforward[1] inhibition by pre-ingestion food cues[5–7], slower feedback inhibition by gut nutrients[33, 34], and even slower feedback inhibition upon restoration of energy balance[1, 33]. HONs directly innervate AGRP neurons[35], and orexin/hypocretin peptides rapidly (secs-mins) activate AGRP neurons *in vitro* [36, 37], implying that arousal stimulates hunger. This simple interpretation creates a paradox. First, it fails to explain why appetitive food cues rapidly activate HONs [38] but inhibit AGRP neurons [5]. Second, it is unclear how this excitatory pathway from arousal to hunger neurons is compatible with suppression of ongoing behavior (such as eating) required for optimal vigilance responses[9–12]. The circuit mechanism that resolves this conflict is unknown.

Here, we resolve this paradox by discovering that the HON-to-AGRP connection operates through a local GABAergic inhibitory microcircuit. Using a combination of *in vivo* calcium recording, optogenetics, and targeted molecular knockdown, we show that HON firing indirectly inhibits AGRP neurons by recruiting a local GABAergic relay. Manipulations of this circuit using targeted neuron ablation while recording pupil as arousal signal demonstrate that it is essential for rapid suppression of AGRP hunger signals in different arousal contexts. These findings reveal a fundamental mechanism for how arousal systems exert context-dependent control over hunger, providing a new framework for how the brain balances competing internal and external demands.

## RESULTS

### Evoked and spontaneous arousals correlate with rapid reductions in hunger drive

To define the real-time dynamics between the arousal and hunger signals, we performed fiber photometry recordings in mice selectively expressing the fluorescent neural activity indicator GCaMP6 in HONs or AGRP neurons, while also tracking pupil diameter (a widely use metric of arousal [12]) (Fig. 1A). We first presented an appetitive food cue. Consistent with previous findings [5, 38], the food cue triggered a rapid increase in HON activity and pupil diameter, and a strong inhibition of AGRP neurons, before any food was consumed (Fig. 1B). To determine if this hunger signal suppression was specific to appetitive cues, we next presented an unexpected sound previously unheard by the mouse (white noise, 120 dB, 1sec). This also robustly increased HON activity and pupil dilation. Surprisingly, these markers of arousal were again coupled with a rapid and significant inhibition of AGRP neurons (Fig.1C). Finally, to determine if the inhibitory arousal-hunger coupling is a fundamental principle of brain state, or limited to discrete external events, we investigated the relationship between AGRP and spontaneous, internally-generated fluctuations in arousal. We found spontaneous pupil dilations to be negatively correlated with AGRP activity, i.e. pupil dilations were accompanied by AGRP inhibition (Fig 1D-F). These data establish that the suppression of AGRP cell activity is a general feature of momentary arousals, occurring for both appetitive and non-appetitive stimuli, as well as for spontaneous arousals.

**Fig. 1.**
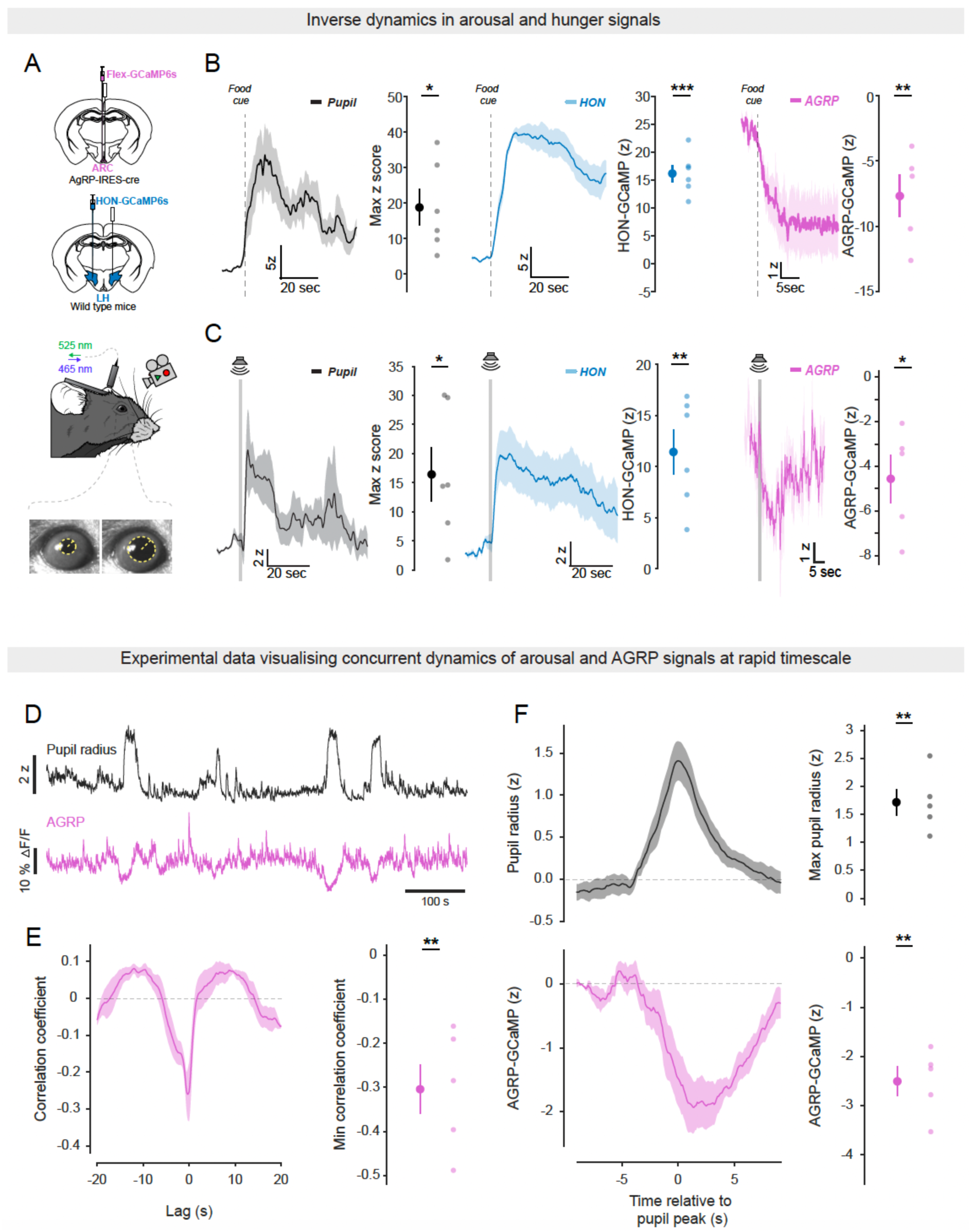
Arousal and hunger signals show opposing dynamics in response to food cue and sudden noise, and this is a fundamental brain state. A. Schematic illustrating the experimental design for calcium imaging and concurrent pupillometry in head-fixed mice. Calcium ARC AGRP neuron activity was recorded in AGRP-IRES-cre mice injected with Flex-GCaMP6s, while calcium LH HON activity was recorded in wild-type mice injected with HON-GCaMP6s. The image below shows the camera setup used for pupillometry, which was performed concurrently with calcium signal acquisition in both experimental groups. B. Dynamics showing the change in HON activity(blue), pupil diameter(black) and AGRP activity(pink) following the presentation of a food cue (vertical dashed line). HON activity and pupil size show a rapid, sustained increase, while AGRP activity exhibits a rapid, sustained suppression. Plots to the right represent the quantified maximal (HONs, pupil) or minimal (AGRP) z-score during the 10s post-cue vs pre-cue baseline (n = 6 for pupil/HONs group, n = 5 for AGRP group, paired t-test, * p = 0.01(pupil), *** p=0.0001(HON), ** p = 0.0088(AGRP)). C. Dynamics showing the change in HON activity(blue), pupil diameter(black) and AGRP activity(pink) following the presentation of a novel sudden noise (vertical dashed line). All three signals show a response: HON and pupil show a transient increase, while AGRP activity shows a transient suppression. Plots to the right represent the quantified maximal (HONs, pupil) or minimal (AGRP) z-score during the 5s post-stimulus vs pre-stimulus baseline (n = 6 for pupil/HONs group, n = 5 for AGRP group, paired t-test, * p = 0.02(pupil), ** p =0.003(HON), * p = 0.01(AGRP)). D. Example recording of pupil radius and AGRP-GCaMP6s activity (representative trace of n = 5 mice). E. Cross-correlation analysis of pupil radius and AGRP-GCaMP6s signal (mean of n = 5 mice). Right, quantification of correlation minima (n = 5 mice, one-sample t-test, ** p = 0.0079). F. Pupil dilation events aligned with corresponding AGRP-GCaMP6s signals (152 events from n = 5 mice). Right, quantification of data, one-sample t-test, ** p = 0.0019 (pupil) and ** p = 0.0011 (AGRP). Throughout the figure, data are expressed as means and shaded areas or vertical bars show s.e.m.

### An intrahypothalamic GABA relay converts HON excitation into hunger cell inhibition

The inhibitory AGRP cell responses to momentary arousals (Fig. 1) cannot be explained by known excitatory inputs from HONs on AGRP cells [35–37]. Therefore, we re-investigated the acute impact of HON activation on AGRP neurons by selectively activating HONs while recording from AGRP neurons. First, to test if HONs can suppress AGRP signals in the absence of pupil input, we selectively optostimulated HONs, while recording the activity of AGRP-GCaMP6s neurons in isolated brain slices (Fig. 2A). HON optostimulation inhibited 50% of AGRP neurons (Fig. 2B, C; the previously-reported excitatory responses were also observed, in 30% of AGRP neurons).

**Fig. 2.**
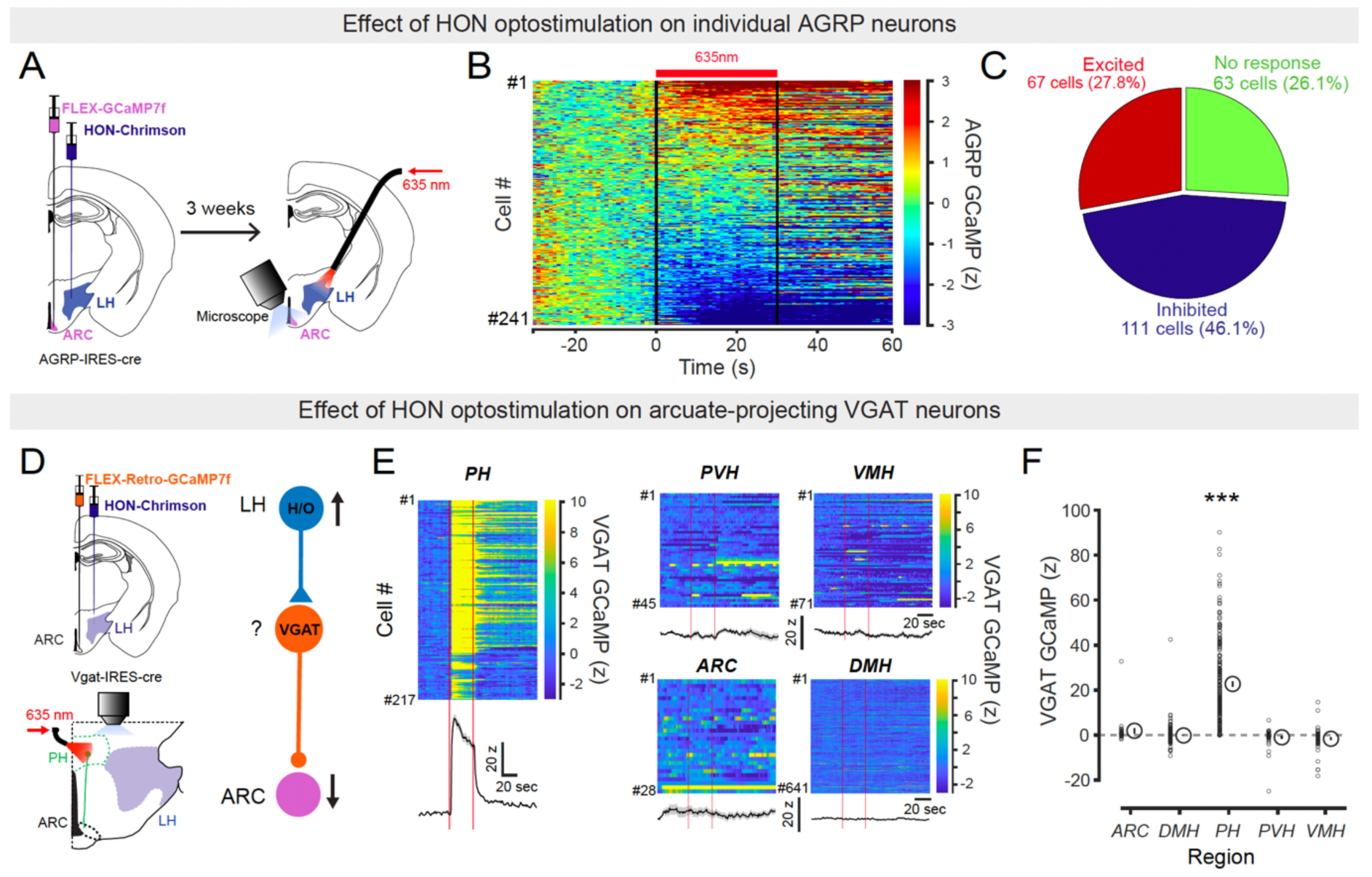
HON optostimulation drives diverse responses in AGRP neurons and region-specific modulation of ARC-projecting VGAT neurons *in vitro*. A. Experimental design for single-cell *in vitro* calcium imaging of AGRP neuron activity and concurrent optogenetic stimulation of HONs. AGRP-IRES-Cre mice were injected with AAV-Flex- GCaMP7f and HON-ChrimsonR in LH. Optogenetic stimulation was provided by 635nm red laser using a fiber optic patch cord aimed at the slice. B. Single-cell AGRP calcium responses, displayed as a heatmap, during red laser stimulation of HONs. The red bar indicates stimulation time. Cells are ranked based on the magnitude of their response (n=241 cells from 3 mice). C. Characterisation of cells identified in (B). The pie chart quantifies the proportion of neurons that were inhibited, excited, or showed no significant response to HON activation (chi-square test, ***p =0.0001). D. Left, experimental design for single-cell *in vitro* calcium imaging of ARC-projecting GCaMP7f in VGAT neurons and stimulation of HON-ChrimsonR axons in various nuclei. Right, diagram of the proposed local circuit showing HON-> ARC projection. E. Heatmaps displaying responses of GCaMP7f-expressing ARC-projecting VGAT neurons to HON ChrimsonR-expressing axon stimulation in different nuclei (ARC, DMH, PH, PVH, VMH). Onset and offset of 635 nm laser stimulation indicated by red lines. Corresponding averaged neuronal responses are shown below each heatmap. F. Quantification of maximal VGAT GCaMP7f response in each region. Note the selective activation of VGAT neurons in PH (n = 214 cells, pre-stimulus baseline vs peak, paired t-test ***p < 0.0001). Statistics for other regions: ARC (p =0.948), DMH (p=0.475), PVH (p =0.131), VMH (p = 0.394). Abbreviations: AGRP, agouti-related peptide; ARC, arcuate nucleus; DMH, dorsomedial hypothalamic nucleus; HON, hypocretin/orexin neuron; LH, lateral hypothalamus; PH, posterior hypothalamic nucleus; PVH, paraventricular hypothalamic nucleus; VGAT, vesicular GABA transporter; VMH, ventromedial hypothalamic nucleus.

We reasoned that this HON-evoked AGRP inhibition may come from arcuate-projecting inhibitory neurons activated by the HON optostimulation, since AGRP neurons do not directly respond to the only inhibitory transmitter made by HONs, dynorphin [39]. To search for such neurons, we targeted the activity indicator GCaMP to inhibitory VGAT neurons innervating the arcuate nucleus (ARC), by injecting Cre-dependent retro-GCaMP into the ARC of VGAT-Cre mice. We then recorded, region-by-region in the slice preparation, the responses of the GCaMP-labelled VGAT→ARC neurons to LH HON optostimulation (Fig. 2D). We found a cluster of HON-activated VGAT→ARC neurons in the mediodorsal posterior hypothalamus (“PH” [40, 41]) but not in neighbouring regions (Fig. 2E, F).

Are these HON-excited VGAT_PH_ neurons involved in HON→AGRP inhibition in vivo? To investigate this, we recorded AGRP-GCaMP activity while optosimulating HONs *in vivo* (Fig. 3A). We found that in awake mice, the HON optostimulation evoked rapid, reversible, and frequency-dependent inhibition of AGRP-GCaMP signals (Fig. 3B, C). This confirms the existence of an HON→AGRP inhibitory pathway *in vivo*. We then repeated these experiments in mice whose PH was injected with VGAT-selective shRNA constructs[42], thus knocking down VGAT expression in VGAT_PH_ neurons (Fig.3D). We found that in these mice, the inhibitory HON→AGRP communication was completely abolished (Fig. 3E, F).

**Fig. 3.**
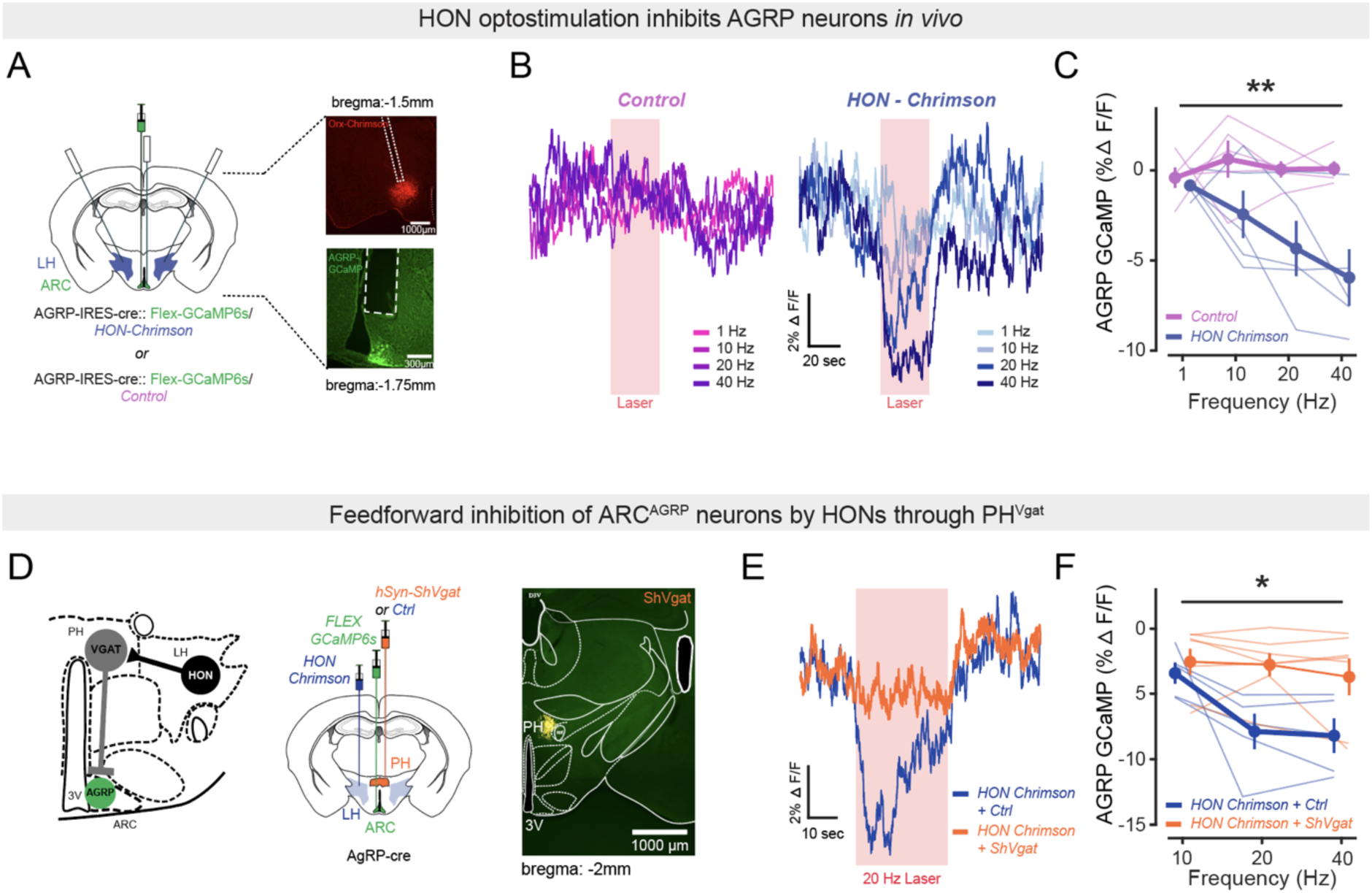
HON optostimulation inhibits AGRP neurons *in vivo*, through a GABA-releasing relay. A. Left, schematic of stimulated (HON) and recorded (AGRP) neurons. Right, examples of corresponding histological verification of expression and fiber placements. B. AGRP-GCaMP6s responses to HON optostimulation (red bar) at different frequencies (means of n = 5 mice in each dataset). C. Quantification of (B) (Two-way repeated ANOVA, **p = 0.0064; n = 5 mice in each group). Post-hoc analysis (ChrimsonR vs. control): 1Hz (p = 0.740), 10Hz (p = 0.053), 20Hz (*p = 0.010), 40Hz (**p = 0.009). D. Left, the proposed feedforward inhibitory circuit: LH^HON^ -> PH ^VGAT^ -> ARC^AGRP^. Middle, experimental design for functional circuit experiment, selective knockdown of VGAT in PH, and simultaneous HON optostimulation and AGRP neuron recording. Right, example image showing viral expression of ShVgat. E. Averaged temporal dynamics of AGRP-GCaMP6s responses to HON optostimulation (laser bar) at 20 Hz in control (means of n =5 mice) and ShVgat (VGAT knockdown in PH) mice (means of n = 7 mice). F. Quantification of (E) (Two-way repeated ANOVA, *p = 0.0166; n = 5 control mice, n = 7 ShVgat mice). Post-hoc analysis (ShVgat vs. control): 10Hz (p = 0.603), 20Hz (**p = 0.004), 40Hz (*p = 0.011). Throughout the figure, data are expressed as means and shaded areas or vertical bars show s.e.m.

Overall, our finding that HON activation leads to AGRP inhibition thus reveals an inhibitory pathway, distinct from the direct excitatory connection between these neurons shown in previous studies. In this pathway, HONs inhibit AGRP neurons through an indirect circuit requiring VGAT_PH_ neurons.

### HON→AGRP indirect inhibition governs appetitive evaluation and adaptive vigilance

We next investigated HON’s role in influencing AGRP cells in three distinct scenarios: appetitive (pre-ingestive) food valuation [5], “what is that?” response to novel stimulus [9], and self-paced pupil dilations that enhace visual scanning of the environment [12, 43].

During appetititve food valuation, we found that HON activation and pupil dilations were greater in response to highly palatable food (peanut butter) than to plain food (chow) (Fig. 4A), similar to the greater inhibition of AGRP neurons by the palatable food vs chow presentation (ref [5], reproduced here in Fig 4D, “control mice”). This indicates that HON and pupil size are normally increased by food cues in proportion to food caloric value. Then, we selectively deleted HONs in adult mice, using HON-specific Diphtheria Toxin Receptor (DTR) cell ablation model[38] (Fig. 4B). In the HON-ablated mice, the inhibitory responses of AGRP neurons to food presentation no longer discriminated between peanut butter and chow (Fig. 4C-E). In contrast, AGRP neuron modulations by internal feedback signals (“hunger hormone” ghrelin, nutrient depletion [5]) were preserved in HON-ablated mice (Supplementary Fig. 1A, B). To also test the necessity of VGAT_PH_ for food value encoding, we also performed these food value experiments in the VGAT_PH_ knockdown mice. Similar to HON-ablated mice, the food value discrimination in AGRP neurons was abolished in the VGAT_PH_ knockdown mice (Fig. 4F, G). Overall, these results demonstrate that HONs are essential for the feedforward[1], sensory prediction of a meal’s caloric value in AGRP neurons, but not for their feedback control by the body’s energy state.

**Fig. 4.**
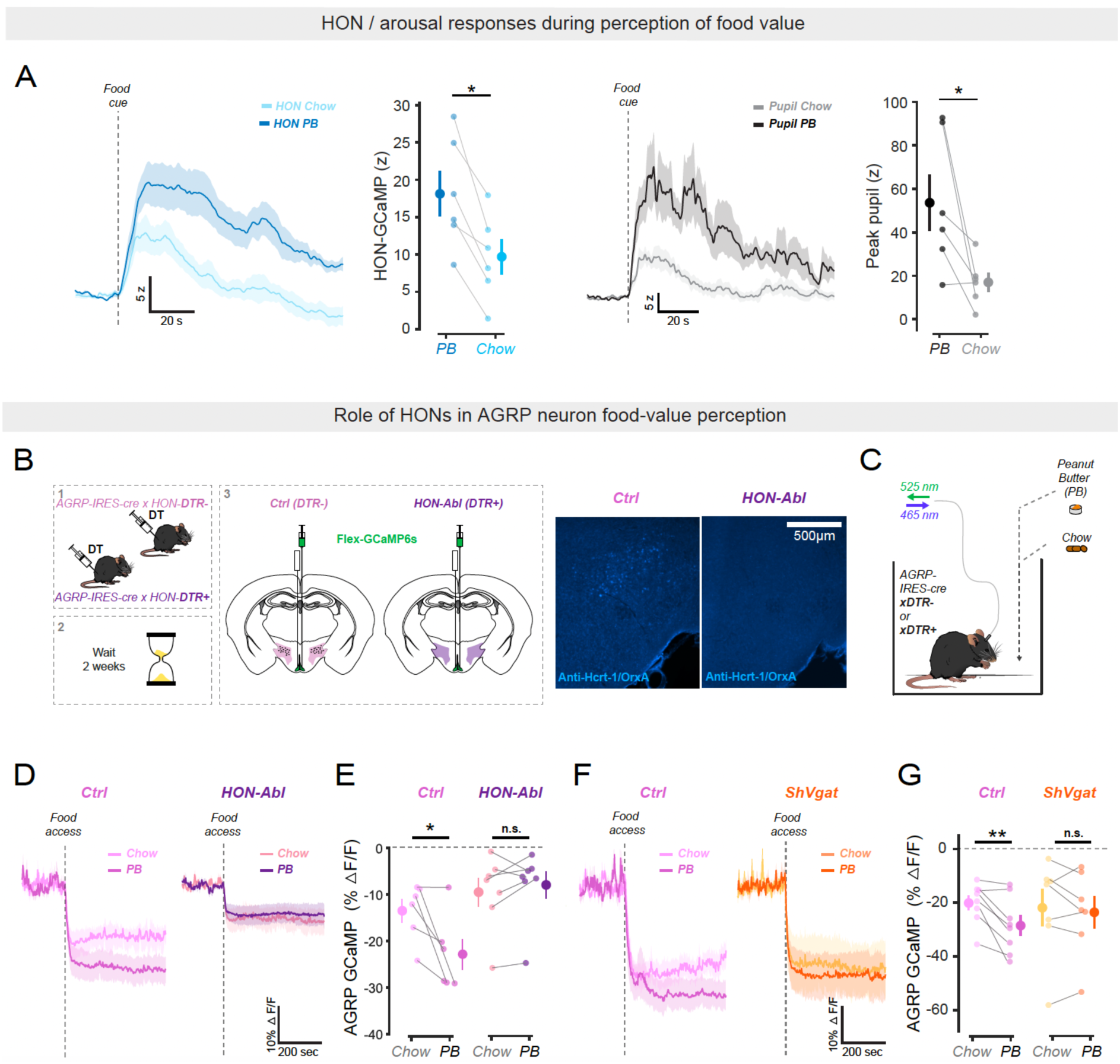
Effect of HONs ablation on rapid AGRP neuron responses to food stimuli. A. Left, HON responses to different food cues (PB = peanut butter). Right, Quantification of HON responses (n =6 mice, paired t-test, *p = 0.0214). Left, pupil responses to different food cues. Right, quantification of pupil responses (n =6 mice, paired t-test, *p = 0.0412). B. Schematic for AGRP neuron recording in HON-ablated mice (HON-Abl) and control mice (Ctrl). Histology shows an example of orexin immunostaining in control mouse (left) and a HON-Ablated mouse (right). C. Schematic of the “food detection” experiment used to obtain data shown in D-E. D. AGRP neuron responses to presentation of peanut butter (PB, darker colours) vs chow (lighter colours) in HON-ablated (HON-Abl) and control (Ctrl) mice (n = 6 Ctrl, n = 7 HON-Abl mice). E. Quantification of data in D. Ctrl: chow vs palatable food *p=0.042; HON-Abl: chow vs palatable food p=0.302. F. AGRP neuron responses to presentation of peanut butter (PB, darker colours) vs chow (lighter colours) in PH-VGAT knockdown (ShVgat) and control (Ctrl) mice (n = 7 ShVgat, n = 8 Control mice). G. Quantification of data in F. Control: chow vs palatable food **p=0.008; ShVgat: chow vs palatable food p=0.487). Throughout the figure, data are expressed as means and shaded areas or vertical bars show s.e.m.

To study “what is that?” responses to arousing non-appetitive stimuli, we established a paradigm where hungry mice were free to consume milkshake from a spout coupled to a lick sensor. An unexpected noise – previously unheard by the mouse - was played while the mouse was eating (Fig. 5A). Mice rapidly reduced their food intake in response to this novel stimulus (Fig. 5B, top), thus establishing a mouse model for adaptive eating reduction. Because we observed rapid, HON activation, and AGRP neuron inhibition response to the noise (Fig. 1C), we investigated whether HONs are essential for the noise-induced eating reduction. We found that the noise-induced eating reduction was abolished in HON-ablated mice (Fig. 5C; AGRP neuron inhibition was also reduced, Fig 5D). Importantly, the pupils of HON-ablated mice still dilated substantially in response to the noise (Supplementary Fig. 2). This suggests that, during the ‘what is that?’ response, arousal is regulated independently from the hunger suppression, and HON ablation selectively abolishes conversion of arousal into hunger suppression, and not the evoked arousal itself.

**Fig. 5.**
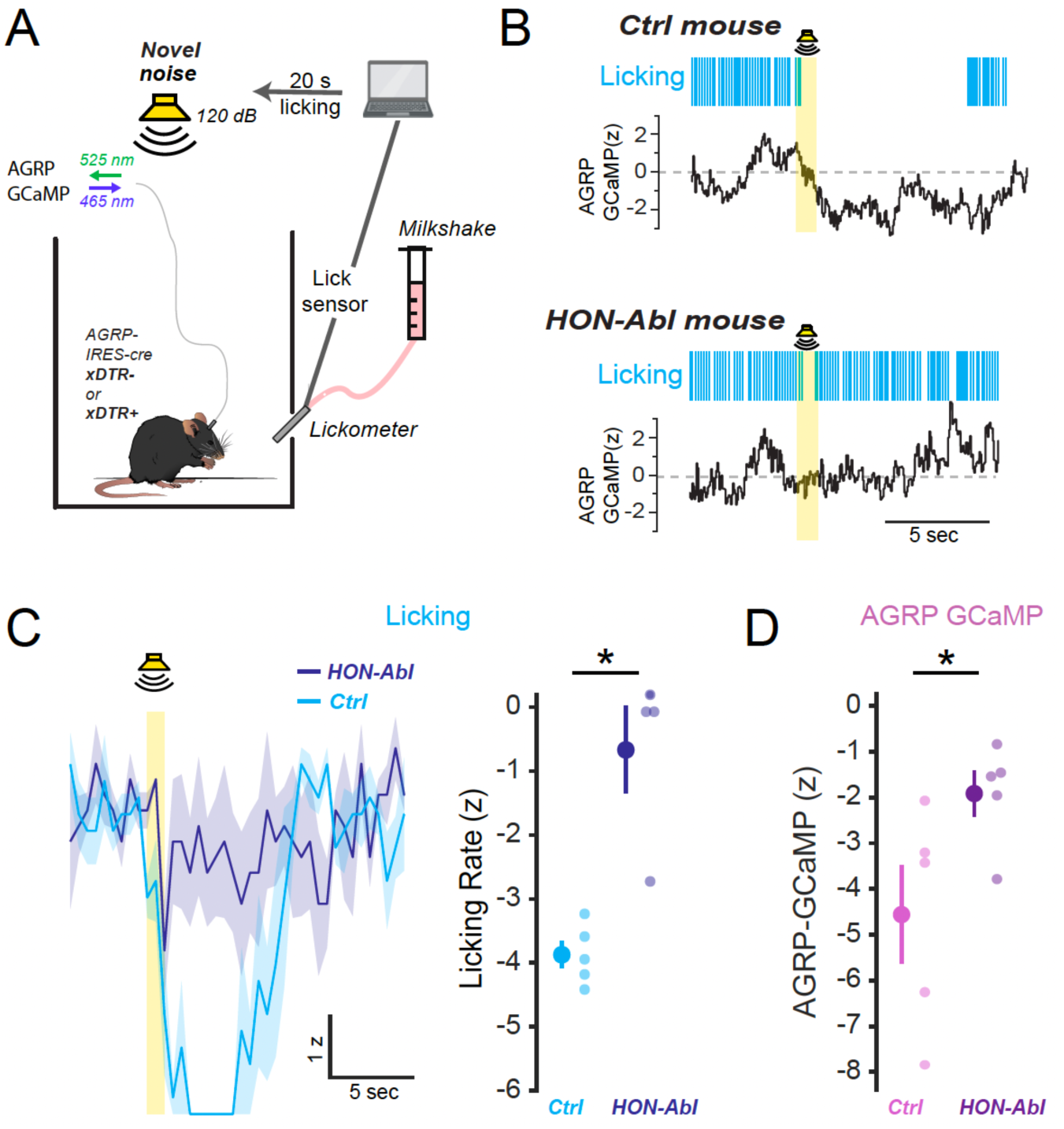
HONs are essential for prioritising adaptive vigilance over hunger. A. Schematic of the “sudden noise” experiment used to obtain data shown in B-D. Following a baseline, mice were given access to milkshake. After 20 seconds of licking, a 1-second novel noise was presented. B. Example effects on eating (top) and AGRP-GCaMP6s (bottom) of the noise (representative trace of Control n = 5 and HON-Abl n = 4 mice) C. Left: Group data for eating responses show in B, mean of Control n = 5, HON-Abl n = 4 mice. Right: Quantification (t-test, **p = 0.0017). D. Quantification of data for AGRP-GCaMP6s response to sudden noise, n = 5 mice per group (one tailed t-test, *p = 0.0279). Throughout the figure, data are expressed as means and shaded areas or vertical bars show s.e.m.

Finally, we found that the inhibitory coupling between self-paced pupil dilations and AGRP cell activity was significantly weakened in the HON-ablated mice (Fig. 6A-C and Supplementary Fig. 3). These findings demonstrate that the HON→AGRP inhibitory circuit does not only react to external events, but also continuously gates the brain’s primary hunger circuit in correlation with the animal’s spontaneous, moment-to-moment arousal state.

**Fig. 6.**
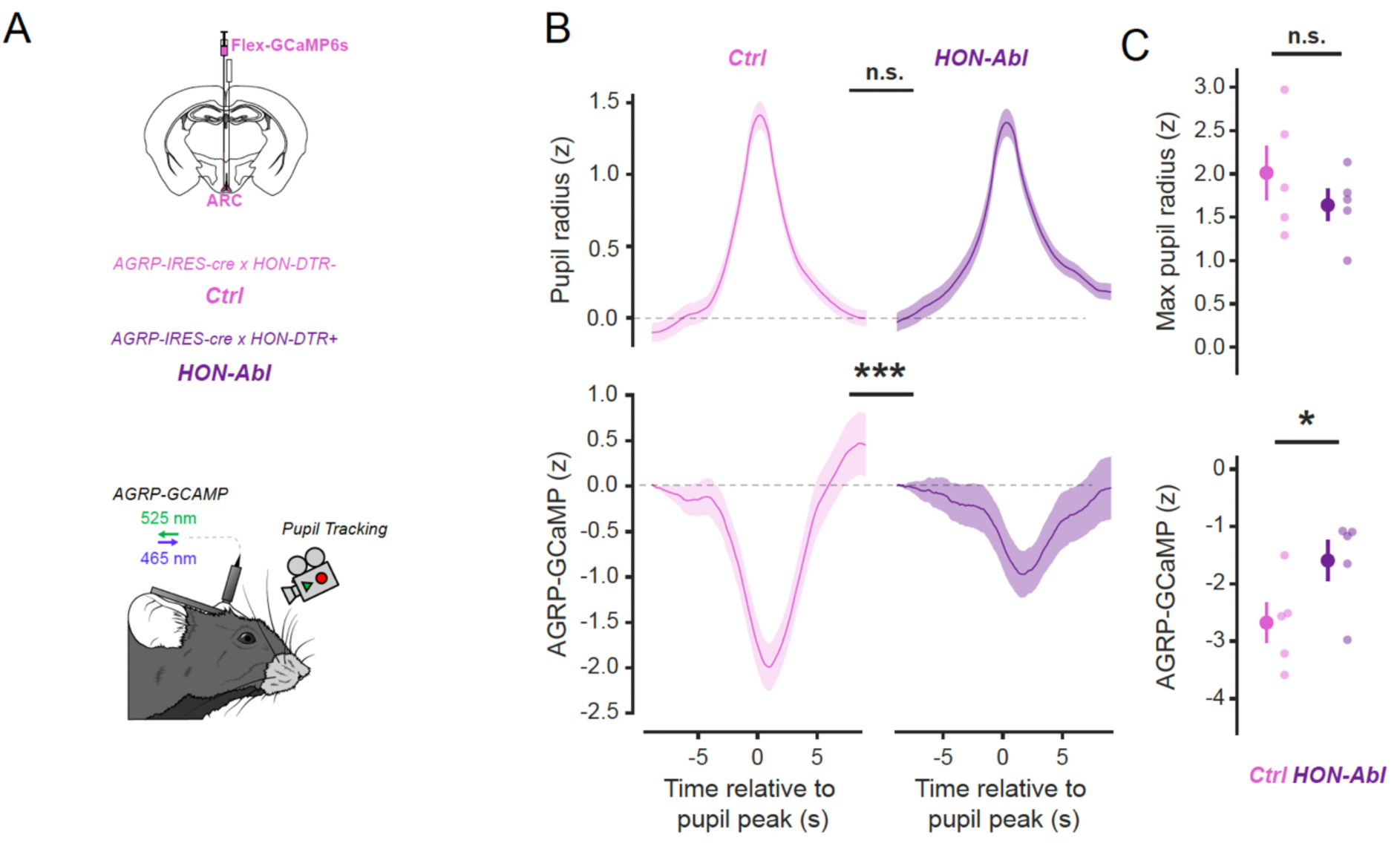
Loss of HONs reduces AGRP inhibition during spontaneous pupil dilations. A. Experimental schematic showing simultaneous AGRP-GCaMP6s recording from the arcuate nucleus (ARC) and pupil tracking in Control (Ctrl) and HON-Ablated (HON-Abl) mice. B. Pupil dilation events (top) aligned with corresponding AGRP-GCaMP6s signals (bottom) in HON-ablated (HON-Abl) and control (Ctrl) mice (196 events from n = 5 control mice, 172 events from n = 5 HON-Abl mice, one-tailed t-test, comparing means of −5 to +5 sec epochs, pupil(top), n.s. = p = 0.590; ***p=0.0003) C. Quantification of data in (B) (n = 5 mice per group). Top: Max pupil radius (one-tailed t-test, n.s. = p= 0.167). Bottom: Min AGRP-GCaMP (one-tailed t-test, *p=0.033). Throughout the figure, data are expressed as means and shaded areas or vertical bars show s.e.m.

## DISCUSSION

A missing part in our understanding of hunger is the role of basic features of brain activity that can be rapidly altered in our daily lives, such as our arousal [12, 44]. Our study addresses this gap by revealing that the brain’s primary hunger signal is under rapid, moment-to-moment inhibitory control from the arousal system. We identified the causal mechanism, uncovering an essential role for HONs and a downstream GABA relay that converts HON activation into AGRP hunger neuron inhibition. Crucially, we demonstrate that this circuit is flexibly deployed to serve two distinct, context-dependent functions: the evaluation of food cues and the rapid prioritization of adaptive vigilance. This discovery reshapes the classic view of hunger control, showing it also dynamically gated by our instantaneous level of arousal to meet immediate behavioral demands (Fig. 7). Below, we propose that this flexible, context-dependent control provides a new framework for understanding value-based decision-making, perception, and symptoms of diseases.

**Fig. 7.**
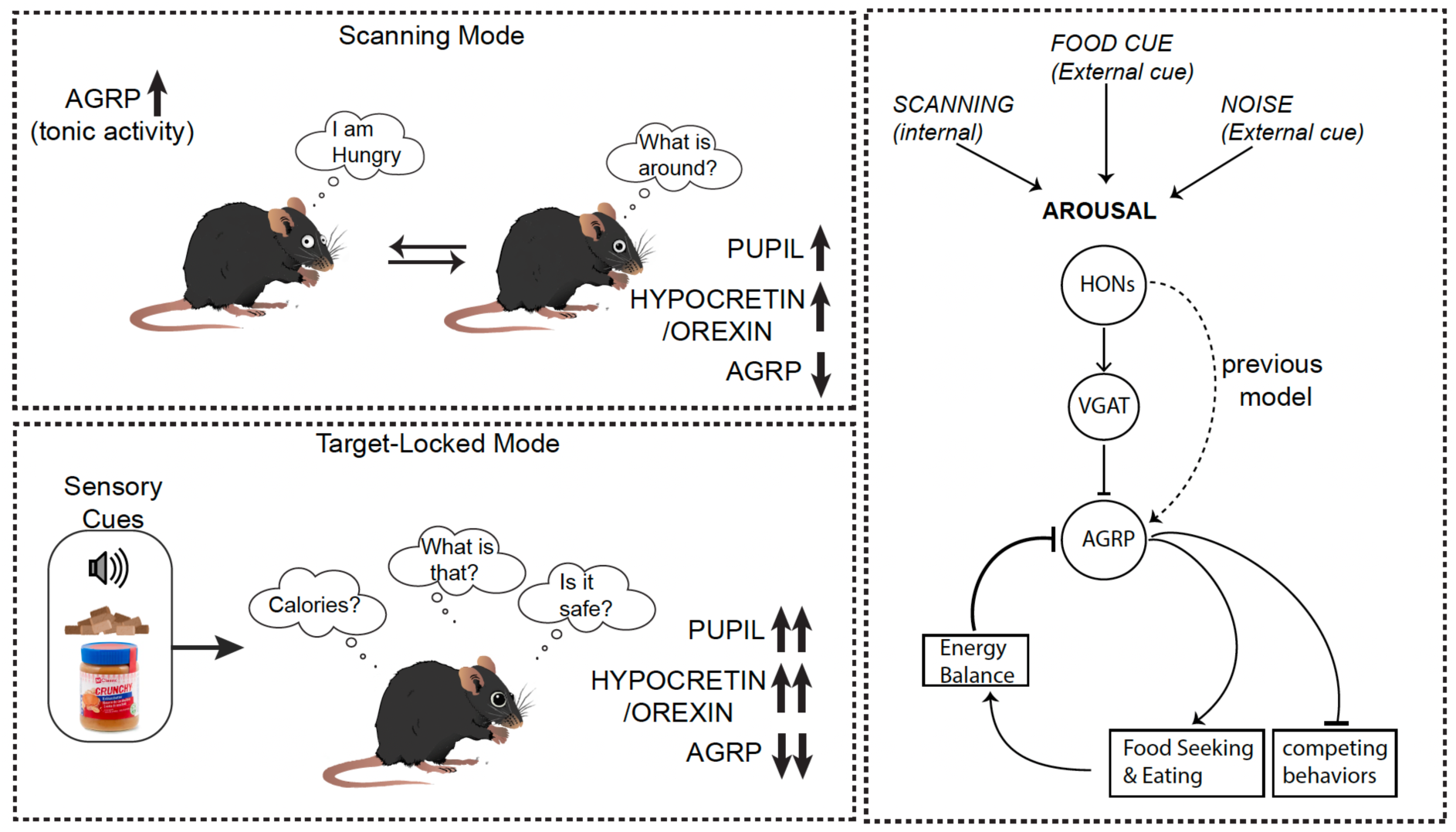
Schematic of proposed significance (left), and relation to previous work (right).

### Physiological implications of arousal-driven hunger suppression for survival

Hunger strongly suppresses other sensations and motivations, which would be maladaptive if an animal cannot be rapidly distracted from hunger to deal with other opportunities or threats [4, 45–48]. We propose that the indirect inhibitory HON→AGRP circuit creates ‘hunger-free’ windows for optimal assessment of opportunities or threats across several important physiological contexts.

In the absence of salient external stimuli, the circuit operates in a proactive, internally-driven scanning mode (Fig. 1D, Fig. 6; summarized in Fig. 7 top left). Here, we found that self-paced pupil dilations coincide with AGRP neuron inhibition and showed that HONs contribute to this co-modulation. This may create windows of sensory acuity [12, 43] while suppressing the distraction of aversive AGRP-driven hunger [7], thus generating dedicated moments of optimal vigilance. Our recordings suggest these vigilance breaks in the hunger drive are frequent and internally-driven. This scanning state can be seen as the brain’s proactive ‘*anything out there?’* mechanism for seeking information.

When an external stimulus is detected, the circuit switches into a reactive, target-locked mode (summarized in Fig. 7 bottom left). External stimuli rapidly activate HONs, increasing multiple aspects of arousal, thus enabling the organism to optimally address dangers or opportunities [15, 16, 24, 26, 49–55]. In an appetitive context, when a food cue is detected, the circuit profoundly inhibits AGRP neurons based on the value of a future meal (Fig. 4). Food cue indicates proximity to a food source, and the more valuable the food, the more likely it is to attract competitors and predators. AGRP suppression in this context may be part of an ‘*is it safe to eat?*’ response ensuring the hungry animal does not over-focus on food, but maintains vigilance for overall survival. Finally, when a strange stimulus is detected, the circuit also strongly inhibits AGRP neurons (Fig. 5), but as part of a global response that suppresses ongoing actions and drives, the classic ‘*what is that?’* response [9–12].

This framework provides a unified mechanism for how a single circuit can produce distinct, context-dependent functions to promote survival by creating rapid breaks from the hunger-induced dulling of other sensations and motivations.

### Relation to previous inconsistencies and pathophysiological phenotypes

It is important to distinguish between rapid and slow influences of arousal on hunger. At long timescales (minutes to hours), being more aroused and awake increases hunger [56]. This is because arousal generally consumes more energy than sleep, and energy depletion is an established cause of hunger. Previous work in vitro suggested a similarly positive relation between rapid manifestations of arousal (HONs) and hunger (AGRP neurons), due to findings that HONs directly innervate and can rapidly activate AGRP neurons [35–37]. This led to models [53, 57, 58] implying that hunger is generated not only slowly after energy depletion, but also rapidly by neural arousal signals. However, this is inconsistent with key clinical observations. In the human disorder, type 1 narcolepsy involves a loss of HONs but increased eating[59, 60] (up to 4 times of normal[61]). In turn, during ageing, there are reports of both an overall decline in food intake (“the anorexia of ageing”[62]) and of difficulties of falling and staying asleep[63, 64], recently linked to ageing-associated hyperexcitation of HONs[65]. Furthermore, in anorexia, there are reports of increased wakefulness and arousal[66, 67].

Our findings reveal an indirect inhibitory pathway from HONs to AGRP neurons which causes arousal and hunger signals to operate in anti-phase. This is more consistent with the clinical observations than the pre-existing models. In states involving loss of HONs (type 1 narcolepsy), we propose that the reported overconsumption in humans [61] and mouse models[38] may be a consequence of losing the HON-dependent “arousal brake” on hunger. Conversely, the hyperexcitation of HONs with aging would be predicted by our model to drive excessive AGRP suppression, thus explaining the anorexia of ageing. Similarly, if the arousal state in anorexia involves elevated HON activity, our model would predict this to suppress the hunger drive, linking a state of anxious arousal directly to appetite loss. Our model thus directly links a single circuit mechanism to key symptoms of distinct human diseases.

### Future directions and outlook

Our findings open important directions for further studies. Hypothalamic recordings and stimulations at single neuron resolution[68] may probe HON subpopulations, providing the indirect inhibition vs the direct excitatory signals to subsets of AGRP cells regulating not only hunger, but also other cognitive and metabolic states [69–71]. While in our experimental settings the inhibitory HON output appeared predominant, this may not be the case in all contexts, and it would be important to investigate the timing and purpose of the excitatory output. More investigations into the relay area, PH [40, 41], are also warranted, such as identifying other regulators of the HON-excited PH neurons, and whether such regulators may gate the indirect signalling from HON to AGRP neurons. Finally, it would be important to investigate additional regulators of hunger downstream of HONs, since our studies do not rule out that their many non-AGRP outputs[23] may regulate feeding, for example, the dopaminergic centers, which also sense and control energy levels and intake [72–76].

More broadly, our findings provide a mechanistic framework for investigating how arousal state influences value-based learning and contributes to the pathophysiology of metabolic and psychiatric disorders. Thus, we establish that a rapid, inhibitory interplay between the brain’s arousal and hunger systems is a fundamental computation for prioritising survival.

## METHODS

### Animal procedures

All procedures followed Swiss Federal Food Safety and Veterinary Office Welfare Ordinance (TSchV 455.1, approved by the Zurich Cantonal Veterinary Office). Adult (older than 22 weeks) female and male C57BL6 mice were used, sex is noted in Supplementary Table 1. Animals were housed in a 12-hour reversed light-dark cycle at 22°C with 55% humidity and had ad libitum access to standard chow (3430 Kliba Nafag, Kaiseraugst, Switzerland) and water unless stated otherwise (Supplementary Table 1). AGRP-IRES-cre mice and VGAT-IRES-cre mice were from Jackson Lab (Jax no. 012899 and 028862, respectively). To study mice without HONs, we selectively ablated HONs using the previously validated hOrx-DTR mouse model[38]. To study AGRP neuron activity in HON-ablated mice, AGRP-IRES-cre and HON-DTR mice were crossed (AGRP-IRES-Cre x hOrx-DTR mice). Experiments were performed during the active (dark) phase of the light cycle.

### Stereotaxic surgeries and viral vectors

Mice were anesthetized with 5% isoflurane, then transferred to the stereotaxic setup (Kopf Instruments, Tujunga, USA) and maintained on 1.5–2% isoflurane. They were given analgesic 30 mins before anaesthesia, and for three days post surgery. Lidocaine was applied to the scalp before an incision was made to access the cranium. Craniotomies were performed using a stereotax-mounted drill (RWD Life Sciences). Viral vectors and implants were positioned at specific coordinates (given below, as mm from Bregma unless stated otherwise). A Nanoject III injector was used to inject the viral vectors at a speed of 1 nl s^−1^.

For AGRP neuron recordings, AAV9.CAG.Flex.GCaMP6s.WPRE.SV40 (titer ≥ 1.9 × 10¹³ vg/mL, from Addgene, MA, USA, 100842-AAV9) was injected unilaterally into the arcuate nucleus of AGRP-IRES-cre, or AGRP-IRES-cre x hOrx-DTR mice, at the following coordinates: anteroposterior −1.75, mediolateral −0.28, dorsoventral −6.1, −6.0 to −5.9 (100nl per site). Optic fibers (400 µm, 0.39 NA, 1.25 mm metal ferrule, Doric) were implanted at dorsoventral −5.85.

For HON recordings, AAV1.hORX.GCaMP6s.hGH (2.5 × 10^12^ GC ml^−1^, Vigene Biosciences), previously demonstrated to selectively target HONs[38], was injected at coordinates: anteroposterior −1.35, mediolateral −/+0.9, dorsoventral −5.7, 5.4 and 5.1 (50nl per site). A GRIN lens (ProView™ Integrated Lens 0.6mm x 7.3mm, Inscopix) was then implanted at dorsoventral −5.1.

For HON optostimulation, injection of AAV9-hORX-ChrimsonR-mCherry (2 × 10^12^ GC ml^−1^, UZH Viral Vector Facility) previously demonstrated to selectively target HONs[77], or control injection (PBS), was performed bilaterally at: anteroposterior −1.45, mediolateral ±2.8, dorsoventral −4.7 and −4.8 (150nl per site). For *in vivo* optostimulation of HONs, optic fibers (200-μm diameter, 0.39 NA, 1.25-mm ceramic ferrule; Thorlabs), angled at a 20-degree angle from vertical, were placed 100 um above injection sites. For *in vitro* optogenetic-assisted probing of HON→AGRP neuron connectivity, AAV1-Syn-FLEX-GCaMP7f (Addgene, titer 1 × 10¹³, 104492) was also injected into the arcuate nucleus (coordinates as above) of AGRP-IRES-cre mice.

For *in vitro* optogenetic-assisted calcium imaging probing functional connectivity from HON to arcuate-projecting VGAT neurons, we injected AAV9-hORX-ChrimsonR-mCherry into the lateral hypothalamus (coordinates as above) and AAVretro-Syn-FLEX-GCaMP7f (Addgene, at titer ≥v7×10¹² vg/mL, catalog# 104492) into the arcuate nucleus (coordinates as above) of Vgat-IRES-cre mice.

For selective knockdown of VGAT expression, we injected VGAT-selective shRNA constructs [36] or PBS as control. The pAAV-hsyn-flex-dsRed-shvgat plasmid (Addgene, Plasmid #67845) was packaged by Charles River as AAV1-hsyn-dsRed-shvgat (titer: 1 × 10¹³ GC/mL). This virus(1:5 dilution) or PBS was injected (150 nl per site) into the posterior hypothalamus (anteroposterior: −2.0, mediolateral: ±0.37, dorsoventral: −4.9) of AGRP-IRES-cre mice. In the same mice, GCaMP6s and hOrx-ChrimsonR viruses were injected, and lenses were placed as described above.

For pupillometry, a custom-made headplate (Protolabs) was cemented to the skull.

All implanted probes and headmounts were secured to the skull with three-part dental cement (Superbond, Sun Medical). Mice were allowed to recover from surgery for at least two weeks before experiments.

### *In vivo* recordings of neural activity

Standard, dual-excitation-wavelength fiber photometry recordings and analyses (described in detail in previous work [52, 78–83] [16, 38, 84–86]) were used to obtain AGRP-GCaMP6s signals. HON-GCaMP6s signals (Fig. 1B,C, Fig.3A) were obtained by whole-field imaging using an INSCOPIX head-mounted microscope, as described in detail in [52]. Fluorescence outputs were converted to normalised photometry signal % ΔF/F as described in [81, 82, 87]. Following the guidelines in [88], for between-subjects visualisations, the % ΔF/F data were then converted to z scores using the function z = (Fn – μ)/σ. Here, Fn is the normalized photometry signal, and μ and σ are the mean and standard deviation (respectively) of Fn of the whole recording session(Fig.1D) or of a local baseline period around an event or a stimulus presentation (Fig.1B,C,F, 2A-F, 3A-F, 4A-G, 5A-E,6C; Supplementary Figs). The local baseline period was −10 to −5 sec in 1F, 6C; and the pre-stimulus baselines are described in remaining supplementary figure legends.

### Combined pupillometry and neural activity recordings

Pupil dynamics in head-fixed mice allowed to run freely on a rotating wheel were recorded and analysed at constant level of light (to avoid effects of light-induced pupil constriction) as previously described[26, 80], using an infrared camera (Blackfly, FLIR Systems, Imaging software Spinnaker SDK, frame rate 30 Hz). Pupil boundaries were tracked using DeepLabCut to identify eight edge points in each video frame. A circle-fitting algorithm in MATLAB calculated pupil radius, enabling precise pupillary measurement across experimental conditions. To compare the response of pupil to noise in control and HON-ablated mice (Supplementary Fig. 2), the subtractive baseline(1s) was calculated as previously described [89].

Photometry and pupil size data were aligned and downsampled to 10 Hz using the function resample, and z-scored across the whole trace. To visualise AGRP-GCaMP6s signals corresponding to similar pupil dilations in different mouse groups, dilation peaks were detected using findpeaks in Matlab (MinPeakProm=1SD) and events in which running was less than (0.05cm/sec) were removed. To analyse temporal correlations between AGRP-GCaMP6s signals and pupil size, AGRP events were detected using findpeaks in Matlab (min peak prominence 5% DF/F), time windows of 10s made around the peak of the event were taken, and cross-correlated with corresponding pupil data using the Matlab function xcorr with ‘coeff’ normalisation.

### *In vivo* optogenetics

Optogenetic stimulation of HONs was performed as described in detail in our previous work [26]. In Fig. 3, 30 sec trains of 5 ms laser pulses at each of the stated frequences were applied at 3 minute intervals. Laser stimulation is repeated 3 times per indicated frequency, in Fig. 3 C,F every line shows the average of responses to these repetitions for each frequency per mouse.

### Brain slice recordings and optogenetics

300 μm hypothalamic coronal slices were acutely dissected and studied as described in detail in our previous work [90]. The slices were placed in a recording chamber of an Olympus microscope, continuously perfused with ACSF[90] at 21°C, gassed with 95% O2 and 5% CO2. To stimulate ChrimsonR expressed in HONs and axons, HEKA Patchmaster software was used to deliver 20 Hz, 30 sec trains of 5 ms pulses of a red laser (635 nm, Laserglow Technologies, 10mW at the fiber tip) through a fiberoptic tip (200 um diameter) placed above mediobasal or mediodorsal hypothalamic areas (defined in Fig. 2, and the coordinates below). Calcium signals were recorded using a UMPlanF l10x objective, Olympus (0.3 NA,10.10mm working distance) and Retiga ELECTRO camera (Imaging). Excitation light for calcium imaging was provided by a 1 Hz 465 nm Sutter Lambda 4DG light source, controlled by HEKA Patchmaster software. Images were acquired using HEKA SmartLUX and HEKA Patchmaster software. Regions of interest (ROIs) were manually labelled using ImageJ, and data were analyzed using MATLAB. Single-cell GCaMP signal was converted to z scores using mean and standard deviation of its baseline (30 sec before laser stimulation). If the z-score exceeded 1.5 SD for at least 5 seconds, the response was categorized as excitatory. Conversely, if the z-score was below −1.5 SD for at least 5 seconds, the response was categorised as inhibitory. Cells with other responses were counted as “No response”. Coordinates for different nuclei were determined using The Mouse Brain in Stereotaxic Coordinates, Franklin & Paxinos, 1997, and are listed in mm from bregma (AP, ML, DV) as follows:

ARC, arcuate nucleus, −1.60 – 1.95, +/− 0.3, −5.9

DMH - dorsomedial hypothalamic nucleus, −1.70-1.95, +/− 0.5, ∼ −5.5

PH - posterior hypothalamic nucleus, −1.85-1.95, −0.4, −4.75

PVH - paraventricular hypothalamic nucleus −0.70-1, +/− 0.20, − 4.75

VMH - ventromedial hypothalamic nucleus −1.60-1.95, +/− 0.5, −5.8

### Food presentation experiments

Animals were habituated to peanut butter (Migros Classic Crunchy, 6.04 kcal g^-1^) and milkshake (energy milk strawberry flavor, 0.76 kcal ml^−1^, Emmi AG) in home cages overnight and to experimental cages for a minimum of two days before experiments started. On experimental days, animals were placed in the experimental cage for 15 min and then different foods (peanut butter, milkshake, or their standard chow [Maintenance Standard diet]) were placed into the cage as previously described [5].

### Lickometer analysis

Palatable food consumption was measured using a custom-built HPF dispenser (cat. no. 161K011, NResearch) with a capacitor-based lick sensor (cat. no. AT42QT-1010, SparkFun Electronics); data were recorded at 500 Hz using custom Python scripts and a digital I/O device (NI-DAQmx, National Instruments). As palatable food, we used milkshake (energy milk strawberry flavor, 0.76 kcal ml^−1^, Emmi AG) because there is abundant evidence that it is highly attractive for both mice and humans [91–93]. 6 µl of HPF was dispensed for every 10 detected licks. In Fig. 5, lick data was z-scored to the local baseline period 5 sec before the sound stimulus.

### Sudden noise experiments

Hungry mice (24 h fasted) were placed in a cage containing the lickometer dispensing milkshake. Mice were previously habituated to both the lickometer and milkshake. After mice licked continuously for 30 sec, a white noise was played through a loudspeaker placed next to the lickometer (1 sec duration, 120 dB), while licking and AGRP neuron responses were recorded (Fig. 5B-D). To quantify the AGRP response to the noise without the confound of licking and moving around the cage, AGRP responses were additionally recorded in the head-fixed setup (as used for pupillometry in Fig. 1C) without food.

### Histology and immunocytochemistry

To validate the locations of AAV virus expression and the fiber optic, we perfused mice with PBS followed by buffered 4% paraformaldehyde (PFA, pH 7.4). Brains were then postfixed in 4% paraformaldehyde for 30h before being placed in 30% sucrose solution for cryoprotection overnight. Brains were then sectioned on a Leica cryostat at 50 μm. For HON immunoreactivity, slices were incubated in primary antibody against hypocretin-A (Abcam ab6214) at a 1:250 dilution in 3% BSA/0.3% Triton solution overnight at 4 °C. Slices were then washed in PBS followed by incubation with AlexaFluor594(at 1:1000 dilution) at room temperature for 2 h. Slices were then washed in PBS. Images were acquired using a fluorescence microscope (Eclipse Ti2, Nikon).

### Data analysis and statistics

Data processing and statistical evaluations were conducted using MATLAB (Mathworks, USA). All animals and data points were included in the analyses, except for those with no signal in fiber photometry at the beginning of experiments. The results of all statistical tests are summarised in figure legends and are fully specified in Supplementary Table 2. T-tests were two-tailed, except where alternative hypotheses were directional in which case one-tailed tests were used (as specified in Supplementary Table 2). P values less than 0.05 were deemed statistically significant. Results and figures are displayed as data mean ± SEM, unless stated otherwise.

## Acknowledgements

This work was funded by ETH Zürich. We thank Nikola Grujic for setting up and advising on pupillometry. We also thank Beate Laube for her help with mouse genotyping. We are grateful to Antoine Adamantidis and Markus Schmid, as well as Brooke Jarvie, Alexander Tesmer and Nikola Grujic, for helpful discussions and suggestions.

## Supplementary Materials

**Supplementary Fig. 1.**
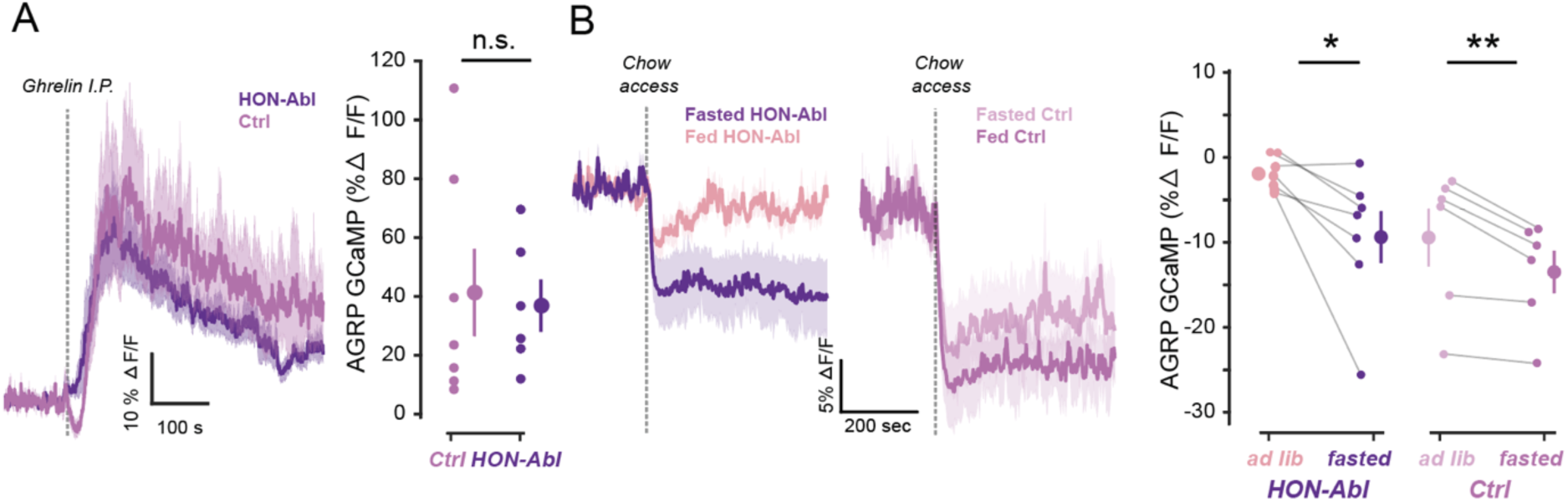
AGRP responses in control vs HON-ablated mice to ghrelin and internal state. A. Left, Comparison of AGRP neuron responses to ghrelin i.p. injection in control vs. HON-Ablated mice. Right, quantification of responses to ghrelin (n = 6 HON-Abl, n = 7 Ctrl; HON-Abl vs. Ctrl, ns = 0.126). B. Left, Average AGRP neuron responses to chow in hungry (24h fasted) or ad lib fed HON-ablated or control mice. Right, Quantification, Fasted vs Fed within each group, paired t=test, *p = 0.03(HON-Abl, n=7 mice), **p = 0.01(ctrl, n=6 mice). Throughout the figure, data are expressed as means and shaded areas or vertical bars show s.e.m.

**Supplementary Fig. 2.**
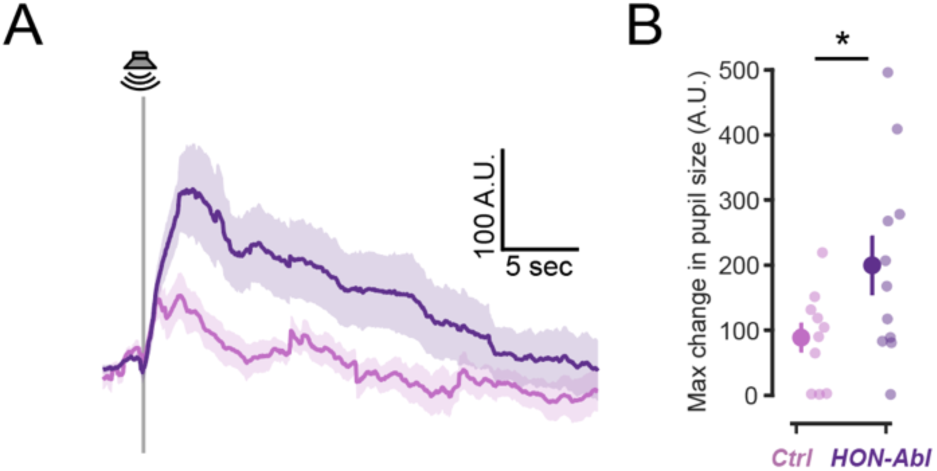
Pupil responses to the sudden noise in HON-ablated mice. A. Temporal dynamics showing the change in relative pupil size following a sudden noise stimulus in fasted control(pink) vs. HON-Ablated mice(purple). B. quantification of peak change in pupil size following the stimulus (n = 11 HON-Abl, n = 10 Ctrl; two-sample t-test, HON-Abl vs. Ctrl, *p = 0.048). Throughout the figure, data are expressed as means and shaded areas or vertical bars show s.e.m.

**Supplementary Fig. 3.**
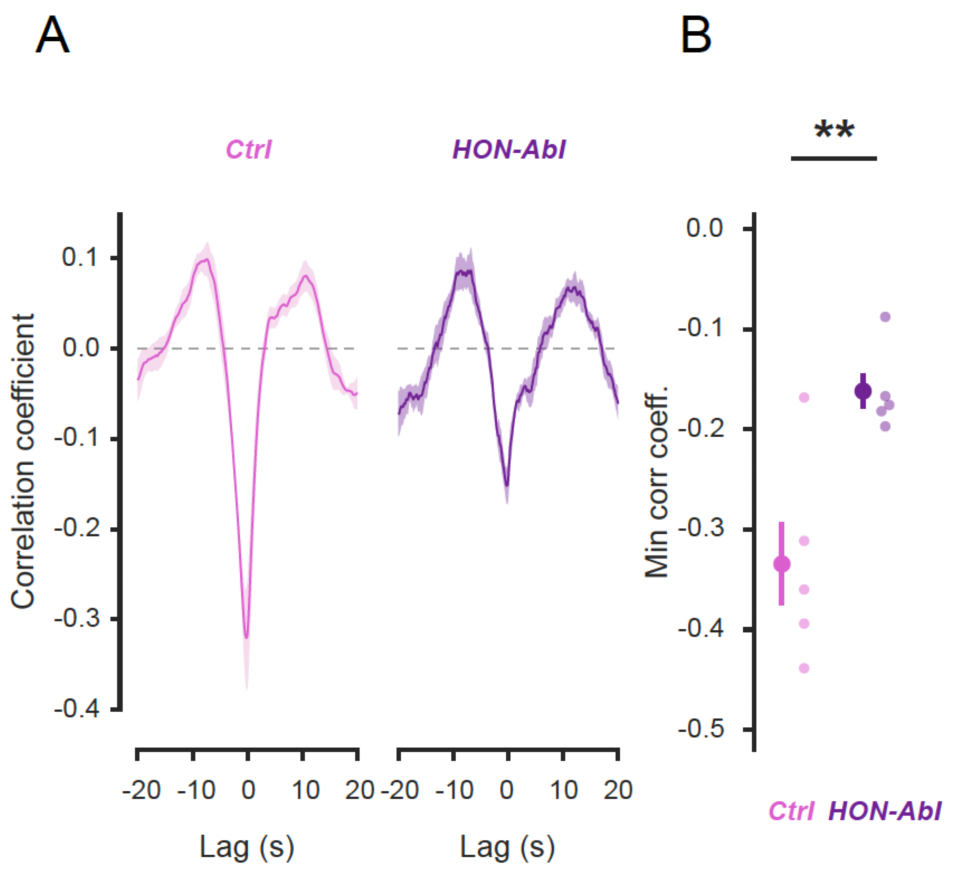
HON ablation reduces correlation of momentary pupil dilations with AGRP neuron activity. A. Cross-correlation analysis of pupil radius and AGRP-GCaMP6s signals (n = 5 mice per group) in HON-ablated (HON-Abl) and control (Ctrl) mice. B. Quantification of correlation minima from (A) (n = 5 mice in each group, one-tailed t-test, ** p = 0.0086). Throughout the figure, data are expressed as means and shaded areas or vertical bars show s.e.m.

**Supplementary Table 1.**
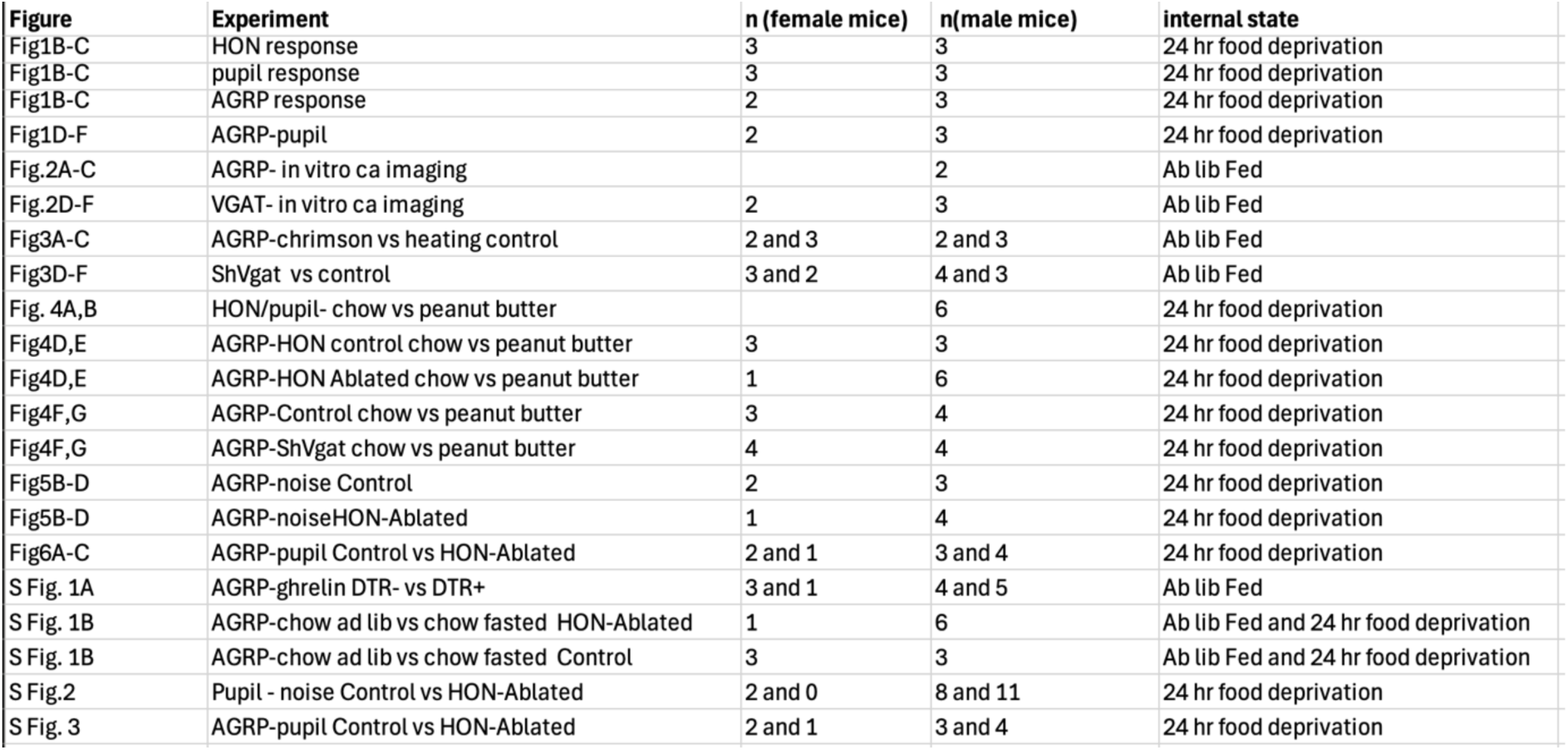
Summary of mouse sexes and nutritive states used in the study. Mice were either ad lib fed or fasted as indicated in the table, for the following resons. H/O neuron optostimulation (Fig. 2) was done in fed mice, to avoid ceiling effects due to fasting-induced activation of H/O neurons. In sleep/wake studies (Fig. 3 and Supplementary 2) mice were fed to avoid effects of fasting on sleep states. In Fig. 4A-F mice were fasted to induce simular continous eating across mice. In Fig. 4G-K, mice were fasted to maximise AGRP neuron responses to food.

| Figure | Experiment | n (female mice) | n(male mice) | Internal state |
| --- | --- | --- | --- | --- |
| Fig1B-C | HON response | 3 | 3 | 24 hr food deprivation |
| Fig1B-C | pupil response | 3 | 3 | 24 hr food deprivation |
| Fig1B-C | AGRP response | 2 | 3 | 24 hr food deprivation |
| Fig1D-F | AGRP-pupil | 2 | 3 | 24 hr food deprivation |
| Fig.2A-C | AGRP- in vitro ca imaging |  | 2 | Ab lib Fed |
| Fig.2D-F | VGAT- in vitro ca imaging | 2 | 3 | Ab lib Fed |
| Fig3A-C | AGRP-chrimson vs heating control | 2 and 3 | 2 and 3 | Ab lib Fed |
| Fig3D-F | ShVgat vs control | 3 and 2 | 4 and 3 | Ab lib Fed |
| Fig. 4A,B | HON/pupil- chow vs peanut butter |  | 6 | 24 hr food deprivation |
| Fig4D,E | AGRP-HON control chow vs peanut butter | 3 | 3 | 24 hr food deprivation |
| Fig4D,E | AGRP-HON Ablated chow vs peanut butter | 1 | 6 | 24 hr food deprivation |
| Fig4F,G | AGRP-Control chow vs peanut butter | 3 | 4 | 24 hr food deprivation |
| Fig4F,G | AGRP-ShVgat chow vs peanut butter | 4 | 4 | 24 hr food deprivation |
| Fig5B-D | AGRP-noise Control | 2 | 3 | 24 hr food deprivation |
| Fig5B-D | AGRP-noiseHON-Ablated | 1 | 4 | 24 hr food deprivation |
| Fig6A-C | AGRP-pupil Control vs HON-Ablated | 2 and 1 | 3 and 4 | 24 hr food deprivation |
| S Fig. 1A | AGRP-ghrelin DTR- vs DTR+ | 3 and 1 | 4 and 5 | Ab lib Fed |
| S Fig. 1B | AGRP-chow ad lib vs chow fasted HON-Ablated | 1 | 6 | Ab lib Fed and 24 hr food deprivation |
| S Fig. 1B | AGRP-chow ad lib vs chow fasted Control | 3 | 3 | Ab lib Fed and 24 hr food deprivation |
| S Fig.2 | Pupil - noise Control vs HON-Ablated | 2 and 0 | 8 and 11 | 24 hr food deprivation |
| S Fig. 3 | AGRP-pupil Control vs HON-Ablated | 2 and 1 | 3 and 4 | 24 hr food deprivation |

**Supplementary Table 2.**
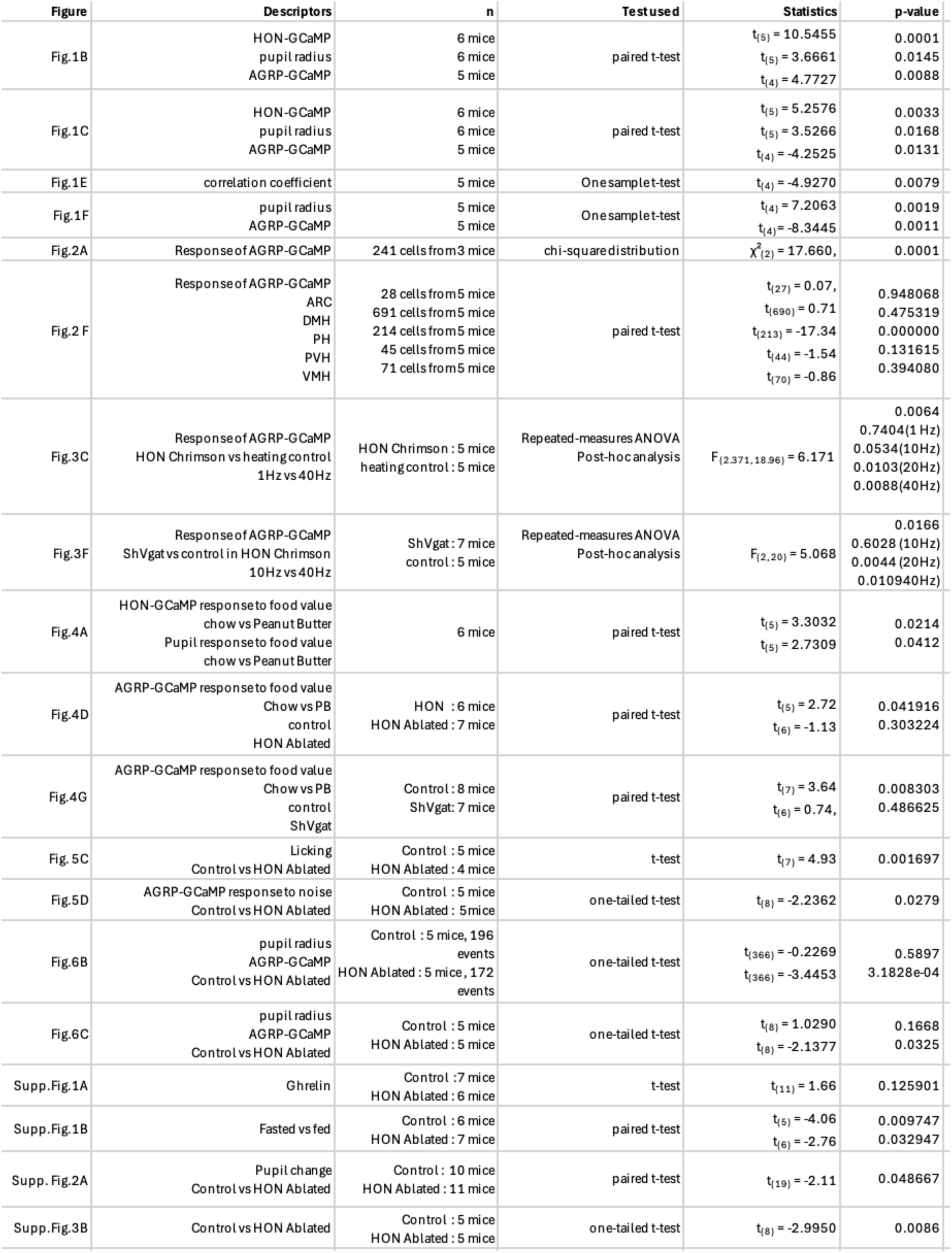
Statistical information.

